# Argonaute bypasses cellular obstacles without hindrance during target search

**DOI:** 10.1101/535575

**Authors:** Tao Ju Cui, Misha Klein, Jorrit W. Hegge, Stanley D. Chandradoss, John van der Oost, Martin Depken, Chirlmin Joo

## Abstract

Argonaute (Ago) proteins are key players in gene regulation in eukaryotes and host defense in prokaryotes. For specific interference, Ago relies on base pairing between small nucleic acid guides and complementary target sequences. To efficiently scan nucleic acid chains for potential targets, Ago must bypass both secondary structures in mRNA and single stranded DNA as well as protein barriers. Through single-molecule FRET, we reveal that lateral diffusion is mediated mainly through protein-nucleic acid interactions, rather than interactions between the guide and targeted strand. This allows Ago to scan for targets with high efficiency but without maintaining tight contact with the DNA backbone. Real-time observations show that Ago “glides” short distances over secondary structures while using intersegmental jumps to reduce scanning redundancy and bypass protein barriers. Our single-molecule method in combination with kinetic analysis may serve as a novel platform to study the effect of sequence on search kinetics for other nucleic acid-guided proteins.

## Introduction

Target recognition by oligonucleotide guides is essential in cellular development, differentiation and immunity^1,2^. Argonaute (Ago) proteins are key mediators of the target interference process, by utilizing short oligo-nucleotides (∼20-30 nt) as guides for finding complementary target sequences^3,4^. The guide-target interaction initiates through Watson-Crick base pairing at the “seed” segment at the 5’ part of the guide, after which target binding propagates downstream along the guide, resulting in target interference^5^.

While eukaryotic Argonautes use RNA guides to target RNA, prokaryotic Agos (pAgo) have been demonstrated to vary with respect to the nature of their guide and target^6-8^. Depending on the pAgo type, it uses either DNA or RNA guides to target single-stranded DNA (ssDNA), RNA or both^2^. The ability of pAgos to cleave ssDNA but not dsDNA suggests a physiological role as a host defence system against single stranded mobile genetic elements^6-8^. Recently, a new family of CRISPR-Cas systems have been discovered in archaea that targets ssDNA but not dsDNA suggesting that these defence systems may be more widespread than previously thought^9^.

The number of potential targets encoded in cellular DNA/RNA is vast^5,10,11^, therefore Ago needs to search long stretches of polymer before finding a canonical target. In a previous biophysical study we suggested that human Argonaute 2 (hAG02) uses lateral diffusion along RNA for target search^12^. Yet, the nature of such a mechanism remains unclear, as lateral diffusion alone would lead to excessive re-sampling of potential target sites and to problems at various roadblocks that are present on the target nucleic acids^13,14^.

For other DNA-binding proteins, such as transcription factors (TFs), a multi-step process termed facilitated diffusion **(Table SI)** has been proposed. Use of such a mechanism would lead to reduced sampling redundancy, and the possibility to circumvent obstructions when TFs search for their targets. After recruiting the DNA non-specifically from solution, the protein diffusively scans only a limited section^15-18^, until it dissociates and returns to solution, after which the protein binds to a new DNA sites to find a specific target. In addition to complete dissociation into solution, intersegmental jumping, where a protein transfers between two spatially close-by segments, has been shown to occur for the DNA binding restriction enzyme EcoRV^19^. The complexity of the target search further increases due to the fact that nucleic acids can be covered with other proteins^20^ and structural elements such as hairpins and plectonemes^21^.

Previous studies have shown that certain DNA/RNA-guided proteins interact with DNA through non-specific electrostatic interactions^22-24^, but none so far have studied the nature of these interactions and their contribution on lateral diffusion. Since these lived^12,22,27-29^, we require a method offering high spatiotemporal resolution. Here we make use of single molecule Förster Resonance Energy Transfer (FRET) to elucidate the mechanism of ssDNA target search by a mesophilic Ago from the bacterium *Clostridium butyricum (CbAgo).* We show that *CbAgo* does not remain in tight contact with the DNA backbone, enabling it to bypass secondary structures along the nucleic-acid chain by “gliding” over them without apparent loss in its ability to recognize its target. After gliding locally, the protein is able to reach distant sites (>100 nt) along the DNA through intersegmental jumps, and then resumes gliding. These different modes of lateral diffusion allow Ago to rapidly search through facilitated diffusion, as well as to bypass substantial obstacles during DNA scanning.

## Results

### Single-molecule kinetics of CbAgo binding

To elucidate the complex target search mechanism, we make use of the high spatial sensitivity of single-molecule FRET (Forster resonance energy transfer). We study a minimal Argonaute complex that consists of *CbAgo,* loaded with a 22 nt DNA guide (small interfering DNA, siDNA) (Hegge, Swarts et al 2019). By using total internal reflection fluorescence (TIRF) microscopy, we recorded the interactions of *CbAgo-* siDNA with target DNA. Target DNA was immobilized on a PEG-coated quartz surface in a microfluidic chamber through biotin-streptavidin conjugation. Guide-loaded *CbAgo* was introduced to the microfluidic chamber by flow. The target was composed of a 3 nucleotide (nt) subseed motif embedded within a poly-thymine sequence and labelled with an acceptor dye (Cy5) **(Figure la).** The guide construct was labelled at nt 9 from the 5’-end with a donor dye (Cy3) **(Figure lb).** A 532-nm laser excitation resulted in donor excitation when the protein loaded with the guide DNA interacted with the target DNA. Once the CbAgo-siDNA complex became bound to the target, the proximity of the donor dye to the acceptor dye on the target resulted in high FRET efficiency. This was followed by a sudden disappearance of the signal, indicating that the complex dissociated from the target and diffused into the free solution. Freely diffusing molecules move too rapidly (∼ μs) in and out of the evanescent field for the current time resolution of the experimental setup (100 ms) and were therefore not recorded. We found that *CbAgo* is not able to target dsDNA directly **(Supplementary Fig. la and Supplementary Fig. lb).** Likewise, when a ssDNA target that lacks complementary to the seed motif of the guide was used, only transient interactions were detected **(Figure lc).** To observe target search that involves intrinsically transient interactions, we determined the optimal target DNA motif for recording binding events. The optimal motif should provide binding events longer than our detection limit of 100 ms, but still lead to dissociation events within the time of our measurement (200 s). To determine the optimal motif, the complementarity between guide and target was incrementally extended from nt 2 to 8 of the guide, showing a gradually increasing dwell time of the Ago-siDNA complex. We found that increasing the number of complementary base pairs above 6 resulted in stable binding beyond the photobleaching time **(Supplementary Fig. lc).** To maintain weak interactions, we continued our experiments using a siDNA with three-base complementarity with the target (nt 2-4) **(Figure Id).** Our estimation of the photobleaching rate (1.4 × 10^−3^s^−1^) was an order of magnitude lower than the dissociation rate (2.7 × 10^−2^ s^1^) **(Supplementary Fig. Id),** indicating that photobleaching does not affect our estimation of the dissociation rate.

### Lateral diffusion of *CbAgo*

It was previously shown that an Ago-guide complex does not directly bind a specific target site from solution, but rather binds non-specifically to random positions along a surfaced-immobilized nucleic acid construct^12^. Such non-specific interactions of *Cb*Ago-siDNA along target DNA are too short-lived to resolve in the absence of a canonical target motif **(Figure lc),** and in the presence of such a motif there was still no lateral diffusion visible **(Figure 1f).** As we were unable to resolve lateral diffusion by *CbAgo* from non-specifically bound regions to the target, we questioned whether the observed stable signal for three complementary base pairs is due to stable binding to the target or contains lateral excursions below our time resolution. In case of the latter, measured apparent dwell times **(Figure 1g)** would consist of the combined dwell times of many target escapes through lateral diffusion, each followed by rapid recapture below the detection limit, before *CbAgo* eventually unbinds from the DNA **(Supplementary Fig. lg).** It is shown (See **Methods Kinetic Modelling)** that such a process of repeated recapture would result in an exponential distribution of apparent dwell times, in accordance with **Figure 1g.**

**Figure 1.**
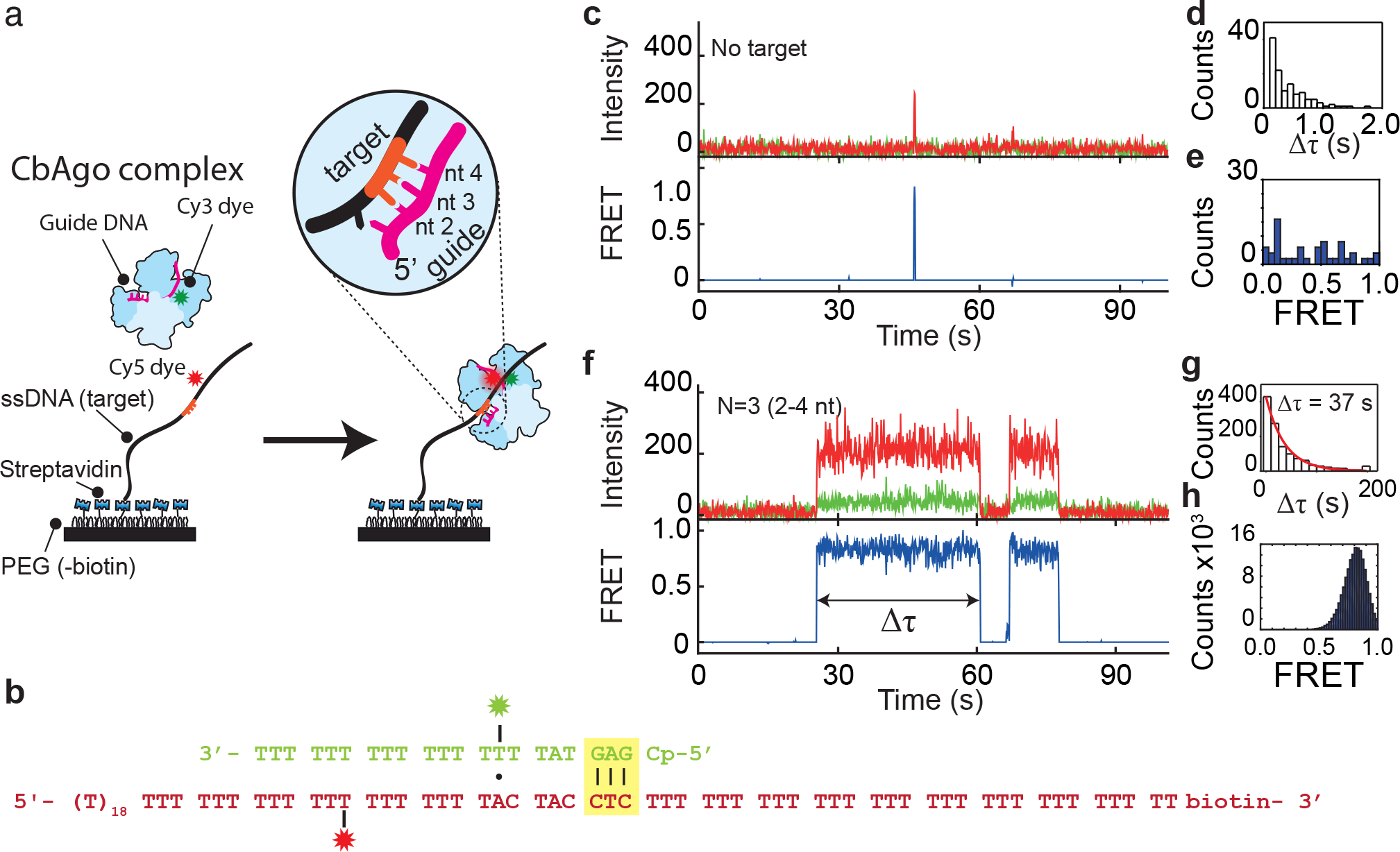
Single molecule imaging of target binding by siDNA:CbAgo complex. **a,** Immobilization scheme of the Argonaute-guide DNA complex. ssDNA is immobilized on a pegylated quartzslide surface. Presence of the Ago-siDNA complex is detected by specific binding to target site (light yellow) resulting in high FRET. **b,** Sequences of guide (green) and target DNA. Guide is labelled on the 9th nucleotide position. **c,** Representative FRET trace of a single molecule experiment at 100mM NaCl showing a transient interaction between CbAgo and a poly-T strand. Time resolution is 100 ms. **d,** Dwell time distribution of the Argonaute in absence of target motif. **e,** FRET values of the transient interactions of d. **f**, Representative FRET trace of a single molecule experiment showing the interaction between CbAgo and a 2-4 nt motif. **g**, Dwell time distribution Dwell time distribution of N=3 binding events with the mean dwelltime of 37 s. h, FRET histogram of binding events, showing a single FRET population for N=3 (2-4 nt) at E=0.78.

To overcome the temporal resolution limit, we adopted a tandem target assay^12,30^. While lateral diffusive excursions from a trap are too short-lived to be resolved in the presence of only a single target, a second target can trap an excursion for long enough to be observed. We placed two identical optimal targets (*N*_*1*_ and *N*_*2*_) separated by 22 nt (**Figure 2a**) along the DNA construct. Both targets base pair only with the first three nucleotides (nt 2-4) of the guide bound by *CbAgo.* As the second target is located further away from the acceptor dye, binding the second target results in a lower FRET efficiency than binding the first target. The respective distance and FRET efficiency between the first binding site (*N*_*1*_) and the acceptor dye (Cy5) remained the same as for the single target assay (E∼0.78), while an additional peak appeared at a lower FRET efficiency for the second target (E∼0.43, **Figure 2d**). This difference in FRET values allows us to determine which of the two sites CfoAgo-siDNA is bound to (**Figure 2b**). After binding to one of the target sites, a majority of the binding events (87.8%) resulted in CbAgo-siDNA shuttling to the other target without loss of FRET signal. Under our standard experimental condition (100 mM NaCl), an average of 13.5 shuttling events occur per binding event (**Figure 2e**). When the experiment was repeated with guides and targets with increased complementary to each other, only 15.1% of the traces showed the shuttling signature within our time window for a 6-nt match (nt 2-7) (**Supplementary Fig. 2f**). This shows that the shuttling signature is controlled by *CbAgo-* ssDNA-motif interactions. With a 6-nt match, the target is so strongly bound and it is less likely that we observe a shuttling event within our observation window.

**Figure 2.**
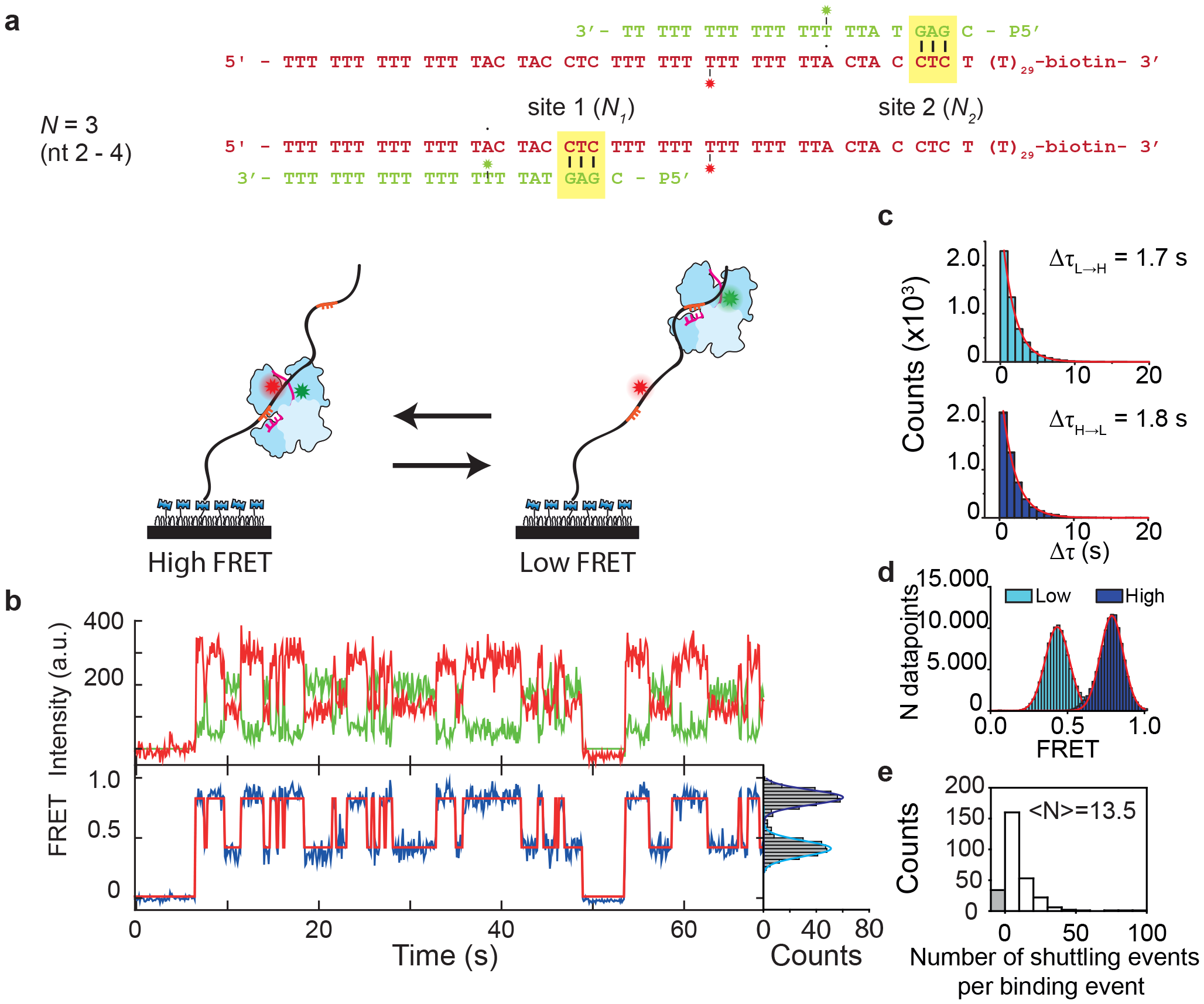
Shuttling signature of CbAgo appears in presence of two targets. **a,** In the top right corner the DNA sequence of guide and target for 22nt separation between targets. Here the distance is defined as the distance from beginning of a target to the beginning of the next target. The placement of the second target (N_2_) results in the appearance of an additional FRET signal, with lower FRET efficiency. **b**, (Top) Representative shuttling trace of a 22 nt separation tandem target at 100 mM NaCl for N=3. (Bottom) The corresponding FRET states (blue) with the fitted HMM trace on top (red). (Right) FRET histogram of the respective time trace. Time resolution is 100 ms. **c**, Dwelltime distributions of respectively the transitions from low FRET state to high FRET state (top) and vice versa (bottom). **d**, FRET histograms of respective states, with peaks at 0.43 and 0.78. **e,** Shuttling event distribution for the same conditions (N=309). Bin size = 10. On average 13.5 shuttling events take place before dissociation. The grey bar (N=33) marks binding events followed by dissociation (no shuttling).

Interestingly, the average dwell time of the first target (**Figure 1g**) decreased from 37 s to 1.7-1.8 s after adding a second target in its vicinity (**Figure 2c**). This observation is in agreement with our lateral diffusion model, since with close-by targets, each sub-resolution diffusive excursion is more likely to be caught in the opposing target.

To further test our claim that the transition between targets occur through lateral diffusion, we extract the time between each shuttling event from traces using single-molecule analysis software^31^. The average time (Δ *τ*_shuttle_) between shuttling events is recorded, the shuttling rate is estimated (*k*_shuttle_ = l/*Δ τ*_shuttle_), and the results are compared to model predictions below.

### Kinetic modelling of lateral diffusion

To determine how lateral diffusion contributes to the shuttling time (the time it takes for Ago to reach the other site), we kinetically model how the this time depends on the distance between traps. The DNA construct is modelled as a series of binding sites along which *CbAgo* will perform an unbiased random walk between neighboring nucleotides. The rate of stepping away from the target is *k*_escape_ in both directions, while at non-specific sites (poly-T) the stepping away is assumed to be near instantaneous—an approximation justified by the fact that lateral excursions are never resolved in the experiments. Hence, the apparent shuttling rate equals the rate at which the protein escapes the initial trap, *k*_*escipe*_, multiplied by the probability to get captured by, shuttle into the other trap (*P*_5_). In the **S.I. (Kinetic Modelling)** we show this probability to scale proportionally with the distance *χ*_Target_ between the targets start nucleotides, resulting in a simple relationship between the average dwell time and the distance between targets.

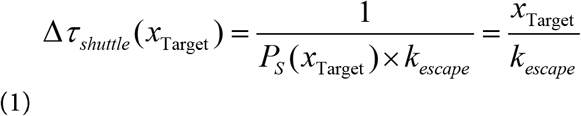

In support of this model, we observed that the apparent shuttling time Δ*τ*_shuttle_(𝓍_Target_) increaseswhen the distance between the targets increases (11, 15, 18 and 22 nt) **(Figure 3**). A fit to **equation (1)** reveals that *Cb*Ago-siDNA complexes escape the target site at a rate of 16 times per second (*k*_escape_ =15.8s^−1^).

**Figure 3.**
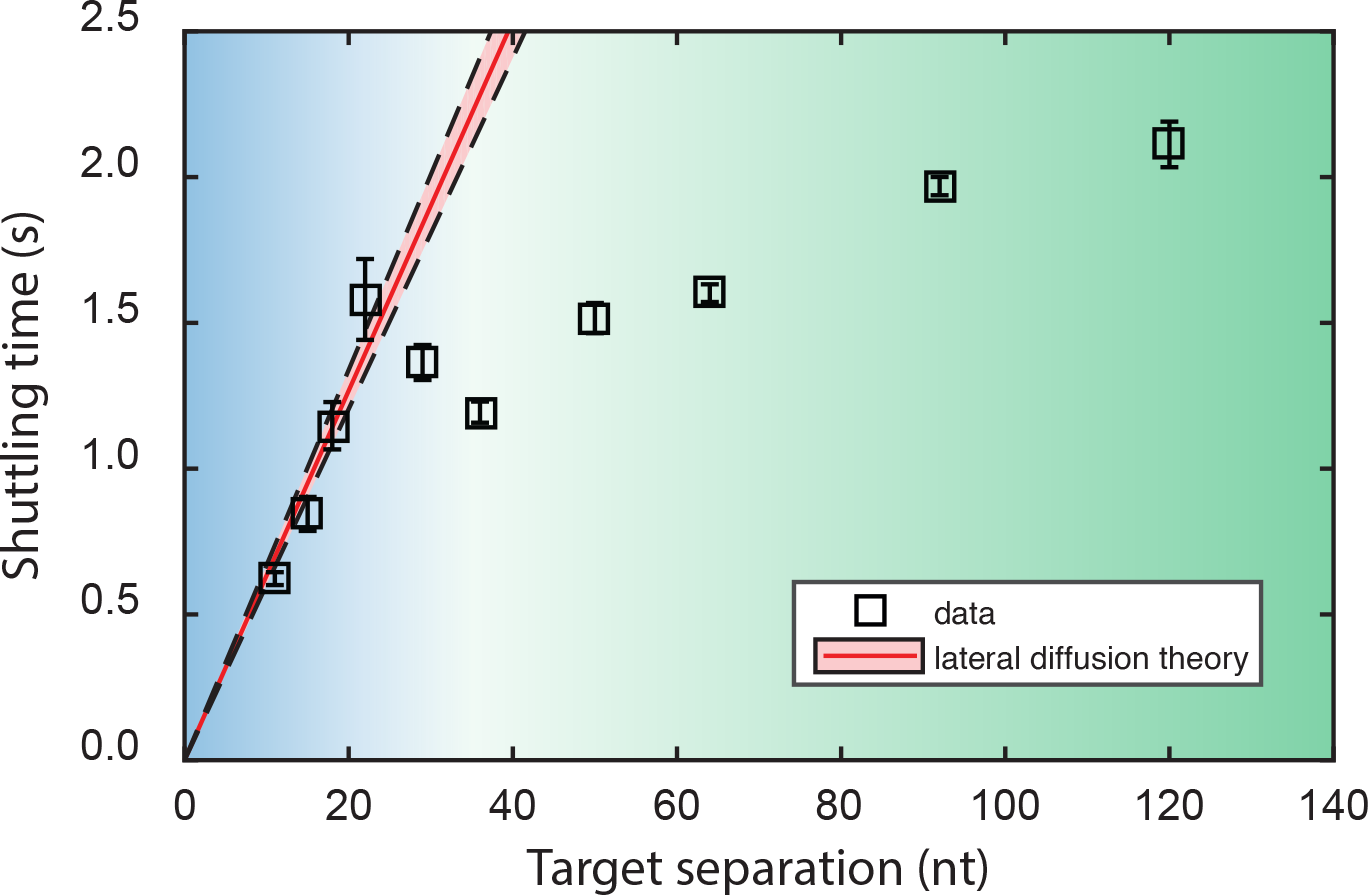
CbAgo shuttling behaviour differs across short and large distances. Shuttling time is plotted versus distance between targets. Squares indicate the mean shuttling time for each DNA construct. The plotted error bars indicate the 95% confidence interval of 10^5^ bootstrapped dwell times. The red line indicates the lateral diffusion model where the first four datapoints are fitted with k_escape_ = 15.8 s^−1^. The shaded red region escape indicates its 95% confidence interval. The blue region indicates where the shuttling time follows lateral diffusion theory. This theory breaks down for larger distances (green).

### Ago probes for targets during lateral diffusion

Next, we placed a third target on the tandem construct **(Figure 4a),** keeping the distance between each set of neighboring targets similar to the target separation in the previous assay (11 nt). We observed three different FRET levels, corresponding to *CbAgo* getting trapped at the three different targets **(Figure 4b).** Using Hidden Markov Modelling (HMM), states can be assigned **(Figure 4b)** and transition probabilities can be extracted **(Figure 4c).** If *CbAgo* returns to solution between binding targets, transitions between any pair of targets will be equally probable, resulting in equal effective rates between all targets. However, if lateral diffusion dominates, transitions between adjacent sites will be favored. The transition probabilities **(Figure 4c)** indicate that most transitions between the two outer targets (from state A to C, or from C to A) proceed through the intermediate target site (state B). The rate to transfer from B to either A or C is greater than that of the opposite path (A or C to B). Using the fitted escape rate from above, *k*_escape_ = 15.8 s^−1^, we predict similar times based on our theoretical model for lateral diffusion **(Figure 4d, S.I. Theoretical Modelling).** With no more free-parameters remaining for this prediction, we take this experimental agreement with our prediction as further evidence of lateral diffusion.

**Figure 4.**
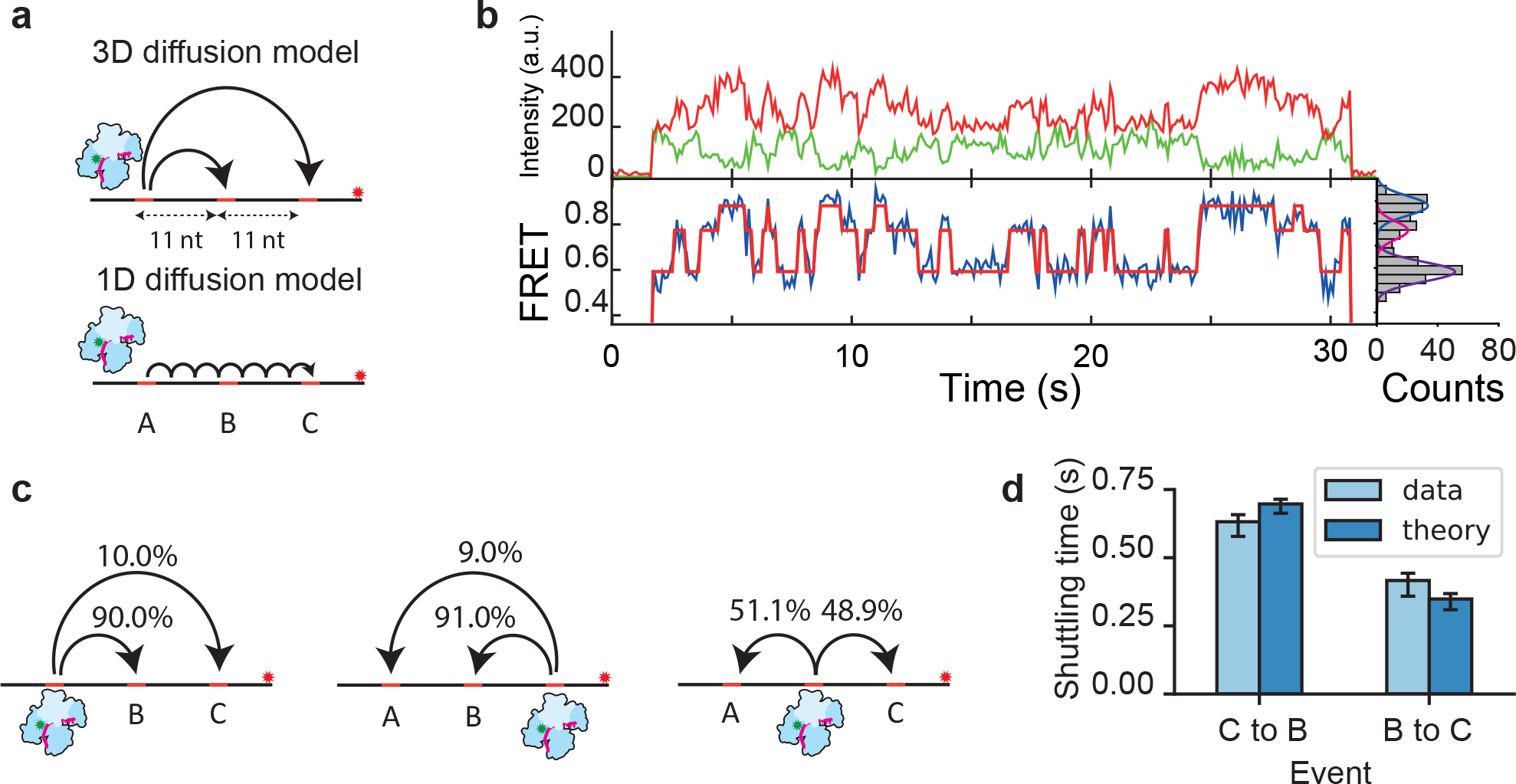
CbAgo undergoes short range diffusion through correlated jumps. **a,** Models for target translocation at short range. In the 3D diffusion model, target dissociation occurs from A followed by random 3D diffusion through solution. In effect, the neighbouring two targets (B and C) will compete for binding. In the lateral diffusion model, the CbAgo complex will have to bypass the adjacent target B before binding to target C. **b,** Representative FRET trace showing the shuttling behaviour between three targets. Top: donor (green) and acceptor (red) intensities. Bottom: FRET trace (blue) and HMM assigned states (red). Right: The fitted states from this data trace. **c,** Transition probabilities from one state to the other two derived from the HMM software. **d,** Experimental values of the shuttling rate of the three target construct were compared against the parameter-free theoretical model that only uses the k_escape_ = 15.8 s^−1^ from **Figure 3**. Error bars indicate the 95% confidence interval acquired from 10^5^ bootstraps.

It is noteworthy that there are about 10% direct transitions from A to C and C to A without any intervening dissociation. The exponential distribution of the dwell times **(Supplementary Fig. 4c**) suggests that at our current time resolution this 10% may be either due to missed events or due to the existence of an additional diffusive mode through which Ago is able to bypass the intermediate target. To conclude, Ago uses lateral diffusion to repeatedly scan DNA segments locally.

### Ago target search is unhindered by structural and protein barriers

Secondary structures are commonly found in mRNA and are also predicted to exist in single stranded viruses^32,33^. It is not known whether *Cb*Ago is able to bypass the numerous junctions it encounters upon scanning a DNA segment. To this end, a Y-fork structure (DNA junction) was introduced as a road block between two targets **(Figure 5a),** while keeping their separation the same as in **Figure 3.** The construct was designed such that the labelled target was partially annealed at the stem with a biotinylated target, thus only annealed constructs were observable on the surface of the microfluidic device. When *Cb*Ago binds to either of the two targets, it can reach the other target only by crossing the junction. Our measurement showed that there was no significant difference in shuttling time between the standard tandem-target construct and the Y-fork construct **(Figure 5b),** indicating that the Y-fork does not impede any of the lateral diffusion modes present. We have previously observed that the CfrAgo-siDNA complex is not able to stably bind to dsDNA, demonstrating that the protein cannot simply go around the junction **(Supplementary Fig. 4c).** Thus, our result suggests that the Ago-siDNA complex does not maintain tight contact with the DNA during lateral diffusion. A weak interaction with the DNA molecule instead allows CbAgo-siDNA to move past the junction.

**Figure 5:**
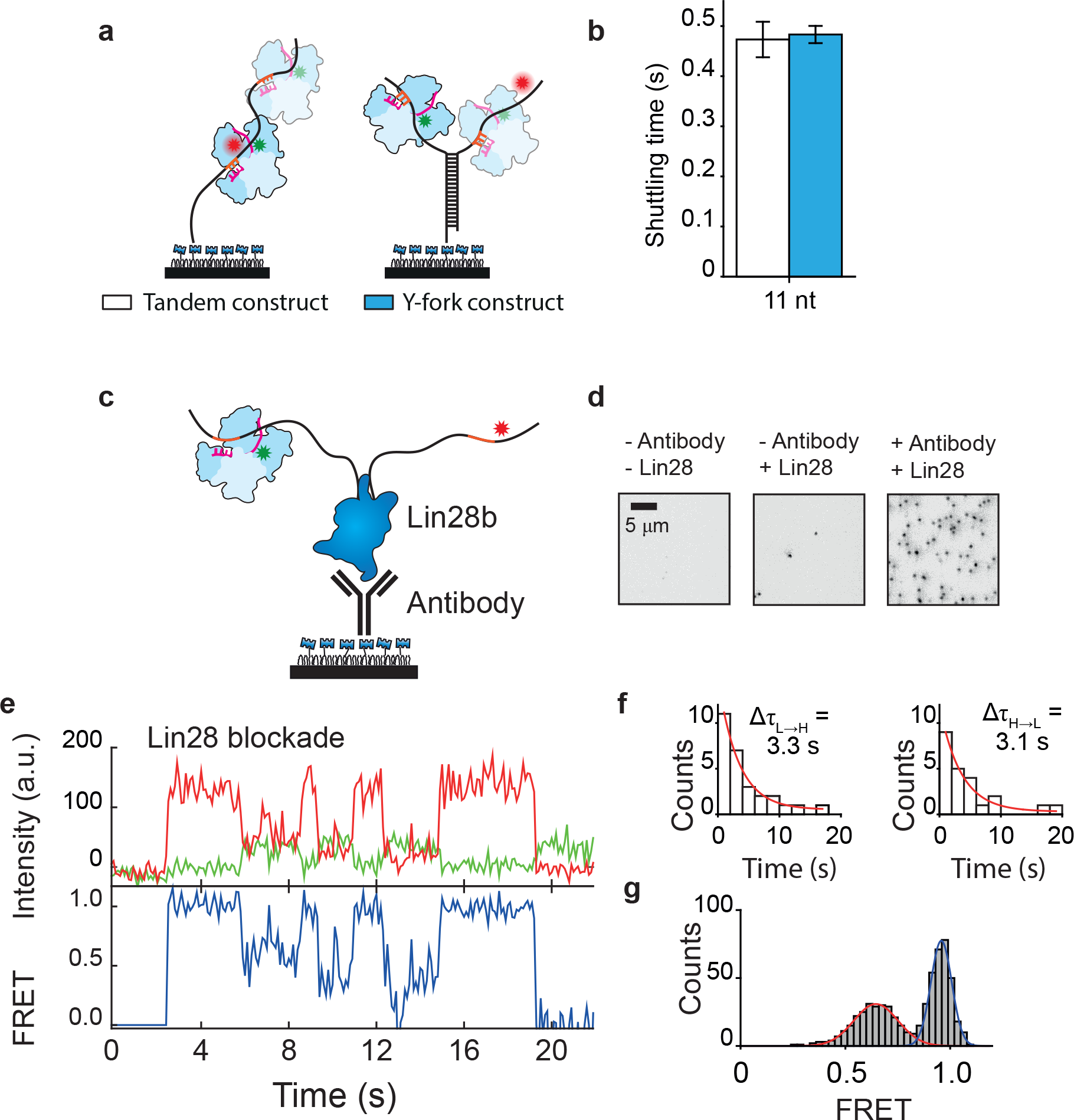
Argonaute can overcome structural and protein barriers. **a,** Schematic drawing of the Y-fork assay (right). CbAgo does not interact readily with the dsDNA junction in the middle so the presence of the junction may interfere with the diffusion. **b,** The shuttling rate of the Y-fork junction (blue bar) compared with the tandem assay (white bar). **c,** Schematic drawing of the his-Lin28b blockade assay. Immobilization happens through a biotin-Anti-His antibody. **d,** An EMCCD image of the acceptor channel. (Left) In absence of Lin28 protein and antibody with Cy5 labeled DNA. (Middle) In absence of antibody, but in presence of Lin28 protein and Cy5 labeled DNA. (Right) In presence of antibody, Lin28 protein and Cy5 labeled DNA. **e,** Example of a shuttling trace with Lin28b located inbetween two targets with an exposure time of 100 ms. f, Individual dwelltimes from low FRET state to high FRET state (left) and vice versa (right). g, FRET histogram (N=46) fit with two Gaussian functions (E=0.64 for the red fit and E=0.95 for the dark blue fit).

Next, we questioned whether *Cb*Ago is also able to overcome larger barriers, such as proteins which are not traversable through lateral diffusion alone. Lin28, a sequence-specific inhibitor of let-7 miRNA biogenesis, has been found to associate sequence specifically to RNA and DNA^34^. His-tagged Lin28 was immobilized on the surface of the microfluidic chamber after which a fluorescent ssDNA fragment was added containing a central Lin28 binding motif and an Ago target motifs on either side **(Figure 5c).** Although the presence of the protein blockade **(Figure 5e)** lowered the shuttling from 0.60 s ^−1^ to 0.27 s^−1^, it did not preclude Ago from reaching the distal site **(Figure 5f).** Since short-range lateral movement is now blocked by the protein barrier, Ago’s ability to move between targets demonstrates that the target search process also allows for intersegmental jumps, in accordance with our observation that the middle target is sometimes skipped when transitioning between the outer targets in **Figure 4d.** Noticeably, the presence of the protein blockade gives rise to FRET fluctuations, broadening the FRET peak **(Figure 5g).**

### Intersegmental jumps allow Ago to rapidly access distant DNA segments

Lateral diffusion is not expected to dominate across large distances, as the linear increase in time of reaching the neighboring (partial) targets **(Equation (1))** would render the search process prohibitively slow. However, when *Cb*Ago was studied with tandem targets that were separated 36 nt or more, we observed that the shuttling still persisted across larger distances **(Figure 3, green region and Table S2).** Together with the evidence of intersegmental jumping above, and the fact that the ssDNA can easily be coiled back to bring the second target close to the Ago protein^35^, we speculate that there is a second mechanism of lateral diffusion: after local scanning for the target through sliding, the *Cb*Ago complex transfers to a different part of the segment that has looped back into proximity of *CbAgo.* This intersegmental jumping mechanism would enable *Cb*Ago to travel to new sites without dissociating, and rescanning of the same segment would be avoided^14,17^.

In order to test this hypothesis, we altered the ionic strength of the buffer solution. Based on the dependence of the single-target off-rate on the ionic strength (**Supplementary Fig. le**), we hypothesized that counterions are not completely expunged when binding to the target sequence. If so, the rate of the intersegmental jumps would also be dependent on salt concentration: increasing the ionic strength should make *Cb*Ago more likely to escape the target motif and more likely to jump to the distant segment. However, at the same time, increased electrostatic screening would also result in a lower binding rate to a nonspecific strand. Depending on which effect is stronger, the shuttling rate may go either up or down when ionic strength is increased. However, if the shuttling took place without disturbing the ionic cloud, the effect from ions would be negligible and therefore the shuttling rate should remain the same.

We used dual-target constructs with 15-nt separation and 64-nt separation **(Figure 6)**, taken from the two different regions in **Figure 3** (indicated by blue and green shading). At a separation of 64 nt, we observed a 13-fold increase of the observed shuttling rate relative to 10 mM NaCl concentration with increasing salt concentration up to 200 mM NaCl. This rate increase suggests that the protein indeed traverses the ion cloud upon long-range shuttling, and that the looseness of association allows for counterions to facilitate dissociation from the target more than it hinders association through screening of the DNA strand.

**Figure 6:**
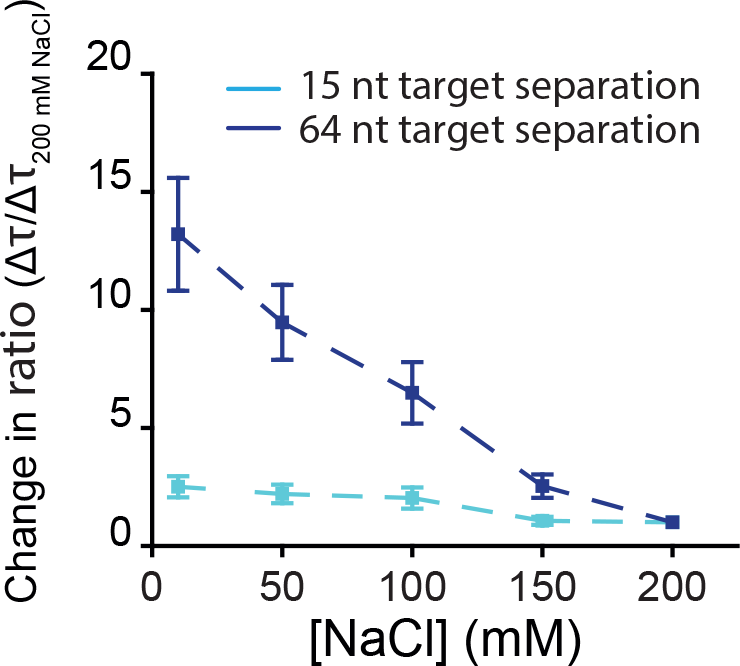
Argonaute target search is characterized by gliding and intersegmental jumps. The relative change in shuttling rate of two constructs from Figure 3, 15 nt separation (light blue) and 64 nt separation (dark blue), normalized against Δτ_shuttling_ at 200 mM NaCl. Error bars indicate the 95% confidence interval of the ratio Δτ/Δτ _10 mM NaCl_. This was determined through 10^5^ times bootstraps.

### Ago uses a gliding mode for local scanning

In contrast, we observed that, for the dual-target construct with 15-nt separation, the shuttling time (when corrected for the dissociation rate) changed roughly only two-fold when the ionic strength of the solution was altered within our range of 10 mM to 200 mM NaCl (**Figure 6**), a modest change compared to 13-fold of the dual-target constructs with 64-nt separation. We hypothesize that during short-ranged lateral diffusion, Ago forces little rearrangements in the ion cloud, and is thus only marginally affected by ionic strength. When targets are close together, this is the main mode of shuttling. Comparatively, for the 64-nt construct, the complex is unlikely to reach the distal site through short-ranged lateral diffusion only, and travels outside the cloud, explaining the strong salt dependence.

In our model, for short distances *Cb*Ago travels predominantly within the ionic cloud, while our Y-fork experiments show that *Cb*Ago need not be in constant physical contact with its substrate during this process (**Figure 4b**). Since neither sliding—characterized by constant binding and complete displacements of counterions, nor micro-dissociations/associations—characterized by longer range jumps^15^-completely capture the properties of this mode of diffusion, we use the term “gliding” to emphasize that it takes short steps, but it does so while being loosely associated with the DNA.

In conclusion, lateral diffusion during *Cb*Ago target search is governed by two distinct modes. For short distances, lateral diffusion takes place through a gliding process characterized by loose contact with the DNA strand while remaining within the ionic cloud. This allows the protein to overcome secondary structures. For larger distances, *Cb*Ago is able to utilize intersegmental jumps to nearby segments.

## Discussion

Within a vast number of potential targets, Ago-guide complexes have to minimize the time spent on non-targets as speed and timing of regulation is crucial for the development of the cell and the host defense^36^. Our single-molecule study shows that Argonaute from C. *butryicum* (*Cb*Ago) uses a target search mechanism distinct from other known DNA binding proteins.

Since the *Cb*Ago searches on ssDNA, not dsDNA, it cannot make use of the structural regularity of double stranded DNA. Bacterial Ago utilizes gliding to scan locally for complementary DNA targets, a mode where the protein is not tightly associated with DNA but stays within the ion cloud. To the best of our knowledge, this mode of lateral diffusion has not been reported for any DNA interacting proteins. In addition, we show that *Cb*Ago is able to move to a new segment via intersegmental jumps, avoiding redundant scanning of the same segment and allowing *Cb*Ago to bypass roadblocks.

In literature, short-range lateral diffusion of target search typically consist either of tight-contact translocation (sliding), or of weakly-bound translocation (hopping)^15^. Our experiments obtained with DNA junctions and protein barriers rule out tight association with DNA strand and thus would suggest that Ago uses hopping (a series of microscopic dissociations and associations) for a dominant mechanism. Unexpectedly, we observed that the weakly interacting short-range lateral diffusion depends only mildly on ionic strength. The absence of a strong ionic strength dependence indicates that, when Ago makes a short exclusion away from a target site, it remains within the ion cloud, only slightly rearranging the ions in its vicinity. Within our experimental conditions, the characteristic Debye distance is estimated to be a couple of nanometers, which is thick enough to accommodate the profile of Ago. Therefore, we propose that Ago is “gliding” through the Debye cloud, displacing some of the counter ions, but not all.

The ability of *Cb*Ago to target specifically ssDNA but not dsDNA **(Supplementary Fig. la and lb)** (Hegge et al, 2019) suggests a role of host defense against mobile genetic elements and ssDNA viruses. In environments where ssDNA viruses can be abundant, such as in sea water, fresh water, sediment, terrestrial, extreme, metazoan-associated and marine microbial mats^37-39^, pAgos targeting ssDNA would be markedly beneficial for the host. Upon entry in the infected cell, ssDNA binding and recombination proteins may associate with the invading nucleic acid, and DNA polymerase will start to generate the second strand. In addition, it is anticipated that secondary structures will be formed in the ssDNA viral genome^32^. This will generate road blocks that may affect scanning by defence systems such as restriction enzymes but not Argonaute. Likewise, insertion of transposons in prokaryotes often proceeds via a ssDNA-intermediate state^40-42^. Here, pAgos may encounter the same obstacles. In case of ssRNA, both in prokaryotes and in eukaryotes, it is well known that complex secondary structures can be formed by base pairing different anti-parallel RNA segments^43-46^. The presence of secondary structures suggest that gliding is necessary for Agos to search along ssRNA. Based on the functional and structural and functional similarities of prokaryotic Agos and eukaryotic Agos^2,12^, we expect eAgo to also glide past RNA secondary structures, minimizing time spent trapped at such structures.

The effect of lateral diffusion on the total target search time is dependent on the roughness of the energy landscape that the DNA binding protein encounters once it binds non-specifically. Theoretical predictions point out that if the roughness exceeds 2 k_B_T, the energy landscape would prevent a protein to diffuse laterally further than a few nucleotides^47^. Here, we expect the variation in the landscape in our poly-T sequence to be minimal for the target search, so that lateral diffusion is able to occur over distances larger than few nucleotides. However, the effect of *in vivo* DNA sequences on the target search remains unexplored, and we expect nucleic acid-guided proteins to encounter a rugged energy landscape during sequence interrogation^48^. We have inferred a 16 s^−1^ escape rate from the 3-nt GAG guide sequence **(Figure 3),** indicating that if a strand were to consists only of GAG in repeating order, the effective diffusion coefficient 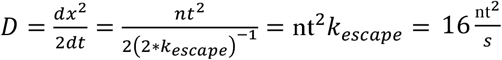. Changing the number of base-paring nucleotides as well as the identity of nucleotides in the guide/target could provide insights into how sequence variation would affect the rate of diffusion for other nucleic acid proteins. Here, we have developed a general platform to could be used to study the effect of sequence identity on search kinetics.

Certain proteins, such as restriction enzymes or DNA repair proteins, are structurally adapted to associate with dsDNA (e.g. the clamp like structure of Msh2-Msh6^49^). However, the currently known structure of pAgos^50,51^ indicates that these proteins lack the capability to firmly embrace and/or unwind dsDNA. This by itself is not surprising, since dsDNA is substantially more conducive for tight interactions than flexible single stranded nucleic acids. Lacking a tight interaction, lateral diffusion for *Cb*Ago is expected to dominate only at short distances, and recurring binding and dissociation events are expected to dominate at long distances. Additionally, since the guide strand only provides the specificity needed for accurate targeting, lateral diffusion could be reliant on the non-specific surface interactions with the protein. We envision that the positive surface charge distribution inside the Ago cleft could orientate the Ago with the guide towards the negatively charged nucleic acid strand **(Supplementary Fig. 4c),** thereby promoting target interrogation while traveling along the target strand. It is unknown whether Ago is able to scan each base during this process or whether it skips over nucleotides. For our triple target construct, we have observed that 90% of the time the middle target traps Ago. It will be of interest to investigate whether this level of effective efficiency of target recognition is achieved by a low trapping efficiency offset by repeated passes over the target before Ago is eventually trapped.

For a longer range target search, we have observed that at 120 nt separation, the shuttling rate remains well above what would be expected for lateral diffusion **(Figure 3).** We hypothesize that coiling of the ssDNA (persistence length ∼ 1 nm) may bring distant segments in close proximity, allowing intersegmental jumps over longer distances (beyond ∼30 nt target separation). Presumably, Ago cannot use intersegmental jumping for covering shorter distances, as noted by the “dip” in **Figure 3,** but the Ago-guide complex might efficiently translocate across longer distances from one place to the other, to bridge sites that are separated >30 nt. Further theoretical modelling is required in order to establish to what extent partitioning different length scales will allow nucleic acid guided proteins to traverse large distances to speed up the search process^47,52,53^.

We hypothesize that similar target search strategies may be used by Agos from different families, which are structurally and functionally similar^2^. For example, in RNA induced transcriptional silencing (RITS), guide-loaded AGOl binds to a transcript after which other proteins are recruited for heterochromatin assembly^54,55^. Similarly, in the piRNA pathway PIWI proteins associate with piRNA in germline cells to bind and cleave transposon transcripts in the cytoplasm^56-58^ or to nascent RNA in the nucleus in order to induce heterochromatin formation^59^. In each of these functions, the reliance on guide-complementary sequential target search likely necessitates the usage of facilitated diffusion strategies to optimize the search time for proper regulation of cell development or gene stability.

## Acknowledgements

We thank Ian MacRae for critical reading and Malwina Szczepaniak, Margreet Docter, Dimitri de Roos, Anna Haagsma and Jan Wignand for technical support. C.J. was supported by Vidi (864.14.002) of the Netherlands Organization for Scientific research. M.K. and M.D. were supported by the Netherlands Organization for Scientific Research, as part of the Frontiers in Nanoscience program. M.D. acknowledges financial support from a TU Delft startup grant. J.v.d.O. was financially supported by two grants from the Netherlands Organization of Scientific Research (NWO; ECHO grant 711.013.002 and NWO-TOP grant 714.015.001).

## Author contributions

T.J.C. S.D.C. and C.J. designed the experiments. T.J.C. performed the measurements. J.H. purified the protein. M.K. and M.D. developed the theoretical model. T.J.C., M.K., J.H., M.D., J.v.d.O. and C.J. wrote the manuscript.

## Competing Interests statement

The authors declare no competing interests.

## Methods

### Purification of CbAgo

The CbAgo gene was codon harmonized for E.coli B121 (DE3) and inserted into a pET-His6 MBP TEV cloning vector (Addgene plasmid # 29656) using ligation independent cloning. The CbAgo protein was expressed in E.coli Bl21(DE3) Rosetta™ 2 (Novagen). Cultures were grown at 37°C in LB medium containing 50μa.g ml-1 kanamycin and 34μg ml-1 chloramphenicol till an ODóOOnm of 0.7 was reached. CbAgo expression was induced by addition of isopropyl β-D-1-ihiogalactopyranosidc (IPTG) to a final concentration of 0.1 mM. During the expression cells were incubated at 18°C for 16 hours with continues shaking. Cells were harvested by centrifugation and lysed, through sonication (Bandelin, Sonopuls. 30% power, 1s on/2s off for 5min) in lysis buffer containing 20mM Tris-HCl pH 7.5, 250mM NaCl, 5mM imidazole, supplemented with a EDTA free protease inhibitor cocktail tablet (Roche). The soluble fraction of the lysate was loaded on a nickel column (HisTrap Hp, GE healthcare). The column was extensively washed with wash buffer containing 20mM Tris-HCl pH 7.5, 250mM NaCl and 30mM imidazole. Bound protein was eluted by increasing the concentration of imidazole in the wash buffer to 250mM. The eluted protein was dialysed at 4°C overnight against 20mM HEPES pH 7.5, 250mM KCl, and 1 mM dithiothreitol (DTT) in the presence of 1 mg TEV protease (expressed and purified according to Tropea et al. 2009^60^) to cleave of the His6-MBP tag. Next the cleaved protein was diluted in 20mM HEPES pH 7.5 to lower the final salt concentration to 125mM KCL. The diluted protein was applied to a heparin column (HiTrap Heparin HP, GE Healthcare), washed with 20mM HEPES pH 7.5, 125mM KCL and eluted with a linear gradient of 0.125-2M KCL. Next, the eluted protein was loaded onto a size exclusion column (Superdex 200 16/600 column, GE Healthcare) and eluted with 20mM HEPES pH 7.5, 500mM KCl and 1mM DTT. Purified CbAgo protein was diluted in size exclusion buffer to a final concentration of 5uM. Aliquots were flash frozen in liquid nitrogen and stored at −80°C.

### Purification of His-tagged Lin28b

The protein was prepared following the protocol of Yeom et al.^61^.

#### Single molecule experimental setup

Single molecule FRET experiments were performed with an inverted microscope (1X73, Olympus) with prism-based total internal reflection. Excitation of the donor dye Cy3 is done by illuminating with a 532nm diode laser (Compass 215M/50mW, Coherent). A 60X water immersion objective (UPLSAP060XW, Olympus) was used for collection of photons from the Cy3 and Cy5 dyes on the surface, after which a 532 nm long pass filter (LDP01-532RU-25, Semrock) blocks the excitation light. A dichroic mirror (635 dcxr, Chroma) separates the fluorescence signal which is then projected onto an EM-CCD camera (iXon Ultra, DU-897U-CS0-#BV, Andor Technology). All experiments were performed at an exposure time of 0.1 s at room temperature (22 ± 0.1 °C)

#### Fluorescent dye labeling of nucleic acid constructs

All DNA constructs were ordered from ELLA Biotech. Nucleic acid constructs that have an internal amino modification were labeled with fluorescent dyes based on the CSHL protocol ^62^.1 uL of 1 mM of DNA/RNA dissolved in MilliQ H20 is added to 5 uL labeling buffer of (freshly prepared) sodiumbicarbonate (84 mg/lOmL, pH 8.5). 1 uL of 20 mM dye (1 mg in 56 uL DMSO) is added and incubated overnight at 4°C in the dark, followed by washing and ethanol precipitation. Concentration of nucleic acid and labeling efficiency was determined with a Nanodrop spectrophotometer.

#### Single molecule chamber preparation

Quartz slides were coated with a polyethylene-glycol through the use of amino-silane chemistry. This is followed by assembly of microfluidic chambers with the use of double sided scotchtape. For a detailed protocol, we refer to ^63^ Further improvement of surface quality occurs through 15 min incubation of T50 and 5% Tween20 ^64^ after which the channel is rinsed with 100 μL T50 buffer. Streptavidin (5 mg/mL) was diluted in T50 to 0.1 mg/mL. 50 μL of this solution is then flowed inside the chamber. This is followed by incubation for 1 min followed by rinsing with approximately 10-fold the volume of the chamber with T50 (10 mM Tris-HCl [pH 8.0], 50mMNaCl). 100pM of DNA/RNA target with biotin construct is then flushed in the chamber, followed by 1 min incubation. This is followed subsequently by rinsing with T50. The chamber is subsequently flushed with CbAgo buffer, containing 50 mM Tris-HCl [pH 8.0], 1 mM Trolox, 1 mM MnCl2, 100 mM NaCl. Guide-loading of apo-CbAGO occurs by incubation of the protein (10 nM) with 1 nM guide construct in a buffer containing 50 mM Tris-HCl [pH 8.0], 1 mM Trolox, 1 mM MnCl2, 100 mM NaCl, 0.8% glucose at 37°C for 30 min. Following incubation, glucose oxidase and catalase is added (0.1 mg/mL glucose oxidase) after which the sample is flushed in the microfluidic chamber containing the DNA targets.

#### Lin28 assay

Immobilization of Lin28b occurred in the following way: 50 μd of streptavidin (0.1 mg/mL) in T50 is flowed inside the chamber and incubated for 1 minute. After this, the chamber is rinsed with approximately 100 μiL of T50. 1 μil of Anti-6X His tag^®^ antibody (Biotin) diluted 100-fold in T50 and subsequently flowed inside the chamber. After 5 minutes, the chamber is rinsed with 100 μL ofT50. Stock of Lin28b (100 μM) is diluted to 100 nM and incubated with the target DNA (10 nM) and 10 mM MgCl_2_for 5 minutes, after which the solution is flushed inside the chamber, followed by incubation of 5 minutes. Lastly, the CbAgo buffer is flushed inside the chamber. Guide-loading of apo-CbAgo occurs in the same way as described above **(Single molecule chamber preparation)** after which the CbAgo:siDNA complex is also flushed inside the chamber.

## QUANTIFICATION AND STATISTICAL ANALYSIS

### Data acquisition and analysis

Fluorescence signals are collected at 0.1-s exposure time unless otherwise specified. For 7-nt target separation, 30-ms exposure time is used. Time traces were subsequently extracted through IDL software using a custom script. Prior to data collection, the location of targets (Cy5 labeled) are found by illuminating the sample with the 637nm laser. Through a mapping file, it subsequently collects the individual intensity hotspots in both the donor and acceptor channel and pairs them up through the mapping file, after which the traces are extracted. During the acquisition of the movie, the green laser is used. Only at the end, the red laser is turned on once more to check for photobleaching of the red dye. Traces containing the fluorescence intensity from the donor and acceptor signal are manually pre-selected occurs through the use of MATLAB (Mathworks), disregarding artefacts caused by non-specific binding, additional binding to neighboring regions and photobleaching.

### Determination of dissociation rate

Binding of Argonaute complex to a single target results in a sudden increase of acceptor signal. The length of these interactions was quantified through a custom script in MATLAB 2015b based on a thresholding algorithm. Briefly, a histogram was made of every data trace, from which the lowest population was fitted with a Gaussian peak. The resulting mean value and standard deviation are then used to distinguish binding events. Intensities that exceeded five times the standard deviation of the baseline (noise) were recognized as a potential binding event. Events that were recognized as potential binding events were marked by the script with a marker for individual checking. Subsequently, the duration of these events were collected and plotted in Origin. Some interactions (at low ionic strength) **(Supplementary Fig. If)** were beyond the observation window of our setup. Hence only a lower limit of the dissociation time could be given. The collected dwell times were bootstrapped through custom code using standard bootstrap algorithms provided by MATLAB. From the resulting distribution, the 95% percentile confidence interval is taken as the error.

### HMM analysis

For assigning states to the FRET traces, a HMM software package is used from Van der Meent et al^31^, which can be found on their github repository (https://ebfret.github.io/). Their software package is optimized for immobilized donor dye molecules on the surface. Here, we immobilize the acceptor dye molecule and hence when no molecule is present, the zero intensity signal in both channels results in large variations in FRET signal, which will result in false positives for the ebFRET software.

Increasing the donor signal and hence artificially creating an extra stable “zero FRET state” is adequate for our purposes, as the distinction between bound and unbound molecules is still made. For the analysis of shuttling traces from constructs where the subseed targets are located far away, the low FRET bound state becomes almost indistinguishable from donor only. Here, this method proves adequate in separating the two states **(Supplementary Fig.** 3).

After assigning states to the collected data, the dwelltimes for low FRET → high FRET and vice versa are extracted. The experimental data shows that there is only one rate-limiting step, in accordance with our theoretical analysis shown below. Using maximum likelihood estimation, the lifetime *Δτ*_*shuttle*_ of the single-exponential distribution 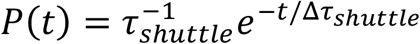 was extracted (the empirical average dwell time equals the ML estimator of *Δτ*_*shuttle*_). The 95% confidence interval was extracted using empirical bootstrapping.

